# Linking photoacclimation responses and microbiome shifts between depth-segregated sibling species of reef corals

**DOI:** 10.1101/2021.10.18.464812

**Authors:** Carlos Prada, Tomás López-Londoño, F. Joseph Pollock, Sofia Roitman, Kim B. Ritchie, Don R. Levitan, Nancy Knowlton, Cheryl Woodley, Roberto Iglesias-Prieto, Mónica Medina

## Abstract

Metazoans host complex communities of microorganisms that include dinoflagellates, fungi, bacteria, archaea, and viruses. Interactions among members of these complex assemblages allow hosts to adjust their physiology and metabolism to cope with environmental variation and occupy different habitats. Here, using reciprocal transplantation across depths, we studied adaptive divergence in the Caribbean corals *Orbicella annularis* and *O. franksi.* When transplanted from deep to shallow, *O. franksi* experienced fast photoacclimation, low mortality, and maintained a consistent bacterial community. In contrast, *O. annularis* experienced higher mortality, and limited photoacclimation when transplanted from shallow to deep. The photophysiological collapse of *O. annularis* in the deep environment was associated with an increased microbiome variability and reduction of some bacterial taxa. Differences in the symbiotic algal community were more pronounced between coral species than between depths. Our study suggests that these sibling species are adapted to distinctive light environments partially driven by the algae photoacclimation capacity and the microbiome robustness, highlighting the importance of niche specialization in symbiotic corals for the maintenance of species diversity. Our findings have implications for the management of these threatened Caribbean corals and the effectiveness of coral reef restoration efforts.

## INTRODUCTION

Understanding how microbial biodiversity interacts with their host’s physiology is essential for understanding animal ecology and evolution (Thompson et al. 2015). Microbial communities often fine-tune their host’s physiology to cope with environmental variation across habitats (Gilbert et al. 2010). Reef-building corals (Cnidaria: Scleractinia) form a symbiotic association with dinoflagellates, which allow corals to thrive on the ocean’s euphotic zone along a strong depth-mediated light gradient (Stoddart 1969). Corals living at different depths possess distinctive physiological and morphological traits to optimize energy acquisition which results from genotypic and phenotypic variation within and between coral species (Vermeij and Bak 2002; Hoogenboom et al. 2008). Coral colonies at different depths may host distinctive symbiotic algae with contrasting photoacclimation capabilities that grant their hosts the ability to thrive in certain light environments (Rowan et al. 1997; Warner et al. 2006). Because of these differences in photoacclimation and the prevalence of specific associations with coral hosts, zonation by light has been regarded as a primary form of niche partitioning in symbiotic corals (Iglesias-Prieto et al. 2004).

While the influence of different species of symbiotic algae on the ecophysiology of reef-building corals has been studied, the effect of other coral-associated microorganisms is less known, specially across depth-segregated species (Rohwer et al. 2002; Pantos et al. 2015). However, the interest on coral-associated microbes and their roles in maintaining health and preventing diseases has increased substantially (Kellogg et al. 2013; Peixoto et al. 2017). From an eco-evolutionary perspective, the evidence suggests that coral-associated bacterial assemblages can be highly variable although “footprints” of unique microbial assemblages appear to be mediated by a combination of host species and local environmental conditions (Rohwer et al. 2002; Marchioro et al. 2020). These patterns indicate that bacterial communities, like photosynthetic dinoflagellates, could also be spatially structured and segregated along environmental gradients.

Recently diverged coral species that differ in their vertical distribution are ideal systems to study the microbiota-animal relationship as a potential basis for habitat specialization. The *Orbicella* species complex, dominant in Caribbean reefs, was initially regarded as one species with ecotypic variation, but recent research revealed three species partially segregated by depth (Weil and Knowlton 1994; Fukami et al. 2004; Levitan et al. 2011). *O. annularis* is a high-light specialist which forms columns with senescent edges, while *O. franksi* is a low-light specialist forming irregular mounds and plates. *O. faveolata* form massive mounds and can overlap with both *O. annularis* and *O. franksi* habitats. The three *Orbicella* species are closely related with incomplete lineage sorting across nuclear and mitochondrial markers (Weil and Knowlton 1994; Fukami et al. 2004). The symbiotic dinoflagellate communities (Rowan et al. 1997; Kemp et al. 2015) as well as the photobiology of this species complex have been extensively studied (Warner et al. 2006; Scheufen et al. 2017), enabling the identification of important differences mediated by environmental gradients. The *Orbicella*-associated bacterial communities have also been examined (Kellogg et al. 2013; Roitman et al. 2020). Therefore, this coral species complex offers an ideal system for the study of how species specialize to live in different habitats through adaptive divergence.

Using a reciprocal transplant experiment between shallow and deep environments in Bocas del Toro (Caribbean Panama), we studied adaptive divergence between the youngest sister species within the *Orbicella* species complex, *O. annularis* and *O. franksi.* We surveyed colonies for survivorship and characterized the algal symbiont and microbial communities across habitats. We also evaluated if these recently diverged species have also diverged physiologically along depth-mediated light gradients. We hypothesize that *O. franksi* and *O. annularis* exploit different light niches, coexisting in Caribbean reefs with minimal competition for space. Our findings suggest that despite being so young (< 500K) (Pandolfi et al. 2002), these two sister species have diverged and fine-tuned their photoacclimation capabilities and microbial symbionts to maximize efficiency in their own light environments.

## MATERIALS AND METHODS

### Reciprocal transplantation

To study the effects of depth and light in *O. annularis* and *O. franksi,* colonies were reciprocally transplanted between shallow and deep environments at Bocas del Toro, Panama (latitude: 9.327222, longitude: −82.203889). The study site is located on the slope of a relative narrow reef protected on all sides by islands and has been monitored for coral spawning for two decades (Levitan et al. 2011). This location is ideal to study adaptation across depths because the vertical distribution of these species is compressed to shallower depths (~2-9 m) compared to other sites in the Caribbean (Van Veghel 1994; Pandolfi and Budd 2008), although maintaining the typical vertical zonation pattern (*O. annularis* in shallow-water and *O. franksi* in deeper-water).

In September 2014, fully pigmented coral clonemate fragments (~5 cm in diameter) were collected from the edges of *O. franksi* colonies and vertically oriented colonies of *O. annularis.* Same genotypes (clonemates) of both species (*n* ≤ 28) were exposed to both shallow and deep environments. Coral fragments were collected from two depths in which each species was abundant: shallow for *O. annularis* (3–4 m) and deep for *O. franksi* (7–8 m). Coral fragments from each species were transplanted to polyvinyl chloride (PVC) panels placed near the original depth of collection (3.5 m and 9.5) where they were left to heal and acclimate for one week. Subsequently, *O. annularis* colonies were transplanted from shallow to shallow (S-S) (*n* = 27) and shallow to deep (S-D) (*n* = 30). Similarly, *O. franksi* colonies were transplanted from deep to shallow (D-S) (*n* = 44) and deep to deep (D-D) (*n* = 28).

To test for differential mortality across depths, we visually inspected colonies six months after transplantation in March 2015. One detached individual from *O. annularis* transplanted deep was discarded from this analysis. A one-tailed Fisher exact test was used to assess differences in survivorship among sites. To standardize the fitness *(i.e.,* survival) advantage on the original depth over the opposite depth for each species, differences in fitness were divided over the average fitness on each particular habitat (Hereford 2009).

Samples were collected in accordance with local regulations under CITES permits PWS2014-AU-002155 and 12US784243/9 and Panama permit number SE/A-94-13.

### Environmental parameters

To characterize the effect of the water optical properties on light availability across depths, we measured the diffuse attenuation coefficient for downwelling irradiance (*K_d_*) at the beginning of the experiment. *K*_d_ was calculated by measuring changes in light intensities across the depth gradient using the cosine-corrected PAR sensor of a Diving-PAM (Walz), previously calibrated against a manufacturer-calibrated quantum sensor (LI-1400, LI-COR). The light intensity at each transplant site, expressed as the percentage of incident light, was calculated (Kirk 2011; López-Londoño et al. 2021). Variation in temperature and relative light levels throughout the duration of the experiment was recorded every 30 min by Onset HOBO data loggers (UA-002-64, Onset Computer Corporation) attached to the PVC panels.

### Photophysiology

To test how depth-dependent light variation affects the photosynthetic condition of corals’ symbiotic algae, we measured the chlorophyll *a* (Chl *a*) fluorescence using pulse amplitude modulated (PAM) fluorometry (Diving-PAM). Measurements were recorded on ten fragments from each species at each depth before transplantation, and every two/three days during the week after transplantation. The effective quantum yield (*ΔF*/*F*_m_’) of photosystem II (PSII) was recorded at noon during peak sunlight exposure, and the maximum quantum yield of PSII (*F*_v_/*F*_m_) at dusk. The maximum excitation pressure over PS II (*Q*_m_) was calculated as *Q*_m_ = 1 - [(*ΔF/F*_m_’)/(*F*_v_*/F*_m_)] (Iglesias-Prieto et al. 2004). *ΔF/F*_m_’ was also recorded *in situ* on coral colonies of *O. annularis* (*n* = 38) and *O. franksi* (*n* = 67) randomly distributed over the full depth range of each species. In order to calculate *Q*_m_ on these colonies, we estimated *F_v_/F_m_* based on a linear regression with data obtained from a sub-sample of colonies randomly distributed over the same depth range (*n* = 10 and *n* = 21 for *O. annularis* and *O. franksi,* respectively). Pearson’s correlation coefficients revealed a strong positive correlation between *F_v_/F_m_* and depth in both *O. annularis* and *O. franksi* (*R*^2^ = 0.85, *p* < 0.01 and *R*^2^ = 0.83, *p* < 0.01), indicating a reliable prediction of *F_v_/F_m_* across depths. We used linear regression models to explore the relationship between *Q*_m_ and depth for *O. annularis* and *O. franksi* based on evidence that *Q*_m_ varies in a pattern that is roughly linear with depth in other coral species (Iglesias-Prieto et al. 2004). An Analysis of Covariance (ANCOVA) was conducted to test for differences in slopes and intercepts among regression models (interaction of species with depth). Due to technical issues with the Diving-PAM (loss in hermeticity), samples from the transplant experiment were transported from the transplant sites to the boat in a dark container to record measurements. During this short period of dark acclimation (<5 min), some components of the non-photochemical quenching could have relaxed (Ralph and Gademann 2005), leading to a slight, yet nearly constant, underestimation of the *ΔF*/*F*_m_’ recorded at noon and, as a result, of *Q*_m_ in all corals. Analyses were conducted using R version 3.6.1 (R Core Team 2015).

### Microbiome

#### Small Subunit Ribosomal RNA (16S) amplicon library preparation and sequencing, sequence quality control and initial data processing

We quantified coral-associated microbiome communities in coral transplants to test if adaptive divergence between *O. annularis* and *O. franksi* is in part due to their microbial communities. Tissue samples were collected at the end of the transplant experiment using 1/8” metal corers by divers wearing Nitrile gloves and were immediately deposited in whirl pack bags. Once returned to the boat, each sample was gently washed with filter-sterile (0.2 μm) seawater, deposited in a sterile cryovial, and immediately preserved in liquid nitrogen. We extracted DNA from coral tissue samples using the MoBio Powersoil DNA Isolation Kit (MoBio Laboratories). Two-stage amplicon PCR was performed on the V4 region of the 16S small subunit prokaryotic rRNA gene (Apprill et al. 2015; Roitman et al. 2020). First, 30 PCR cycles were performed using 515F and 806R primers (underlined) with linker sequences at the 5’ ends: 515F_link (5’-ACA CTG ACG ACA TGG TTC TAC A<u>GT GCC AGC MGC CGC GGT AA</u>-3’) and 806Rb_link (5’-TAC GGT AGC AGA GAC TTG GTC T<u>GG ACT ACH VGG GTW TCT AAT</u>-3’). Each 20 μL PCR reaction was prepared with 9 μL 5Prime HotMaster Mix (VWR International), 1 μL forward primer (10 μM), 1 μL reverse primer (10 μM), 1 μL template DNA (~20 ng/ μL), and 8 μL PCR-grade water. PCR amplifications consisted of a 3 min denaturation at 94 °C; 30 cycles of 45 s at 94 °C, 60 s at 50 °C and 90 s at 72 °C; and 10 min at 72 °C. Amplicons were barcoded with Fluidigm barcoded Illumina primers (8 cycles) and pooled in equal concentrations for sequencing. The amplicon pool was purified with AMPure XP beads and sequenced on the Illumina MiSeq sequencing platform at the DNA Services Facility at the University of Illinois at Chicago. Sequences were submitted to the National Center for Biotechnology Information (NCBI) Short Read Archive (SRA) under project number PRJNA717688.

Initial processing of 16S libraries was performed using the Quantitative Insights Into Microbial Ecology (QIIME; v1.9) package (Caporaso et al. 2010b). Primer sequences were trimmed, paired-end reads merged, and QIIME’s default quality-control parameters were used to split libraries among samples. Chimeras were removed and 97%-similarity OTUs picked using USEARCH 7.0 (Edgar 2010), QIIME’s subsampled open-reference OTU-picking protocol (Rideout et al. 2014), and the 97% GreenGenes 13_8 reference database (McDonald et al. 2012). Taxonomy was assigned using UCLUST and reads were aligned against the GreenGenes database using PyNAST (Caporaso et al. 2010a). FastTreeMP (Price et al. 2010) was used to create a bacterial phylogeny with constraints defined by the GreenGenes reference phylogeny. OTUs classified as “unknown” (*i.e.*, sequences not classified at the kingdom level), chloroplast, mitochondria, or other potential contaminants were removed. Low coverage samples (< 223 useable reads) were omitted. Unless otherwise stated, downstream microbiome analyses and figure generation were performed in R version 3.2.5 (R Core Team 2015) using the phyloseq and ggplot2 packages (Wickham 2009; McMurdie and Holmes 2013).

#### β-diversity group significance and differential abundance testing

To quantify differences among treatments, we used weighted UniFrac (wUniFrac) dissimilarity matrices using OTU-level relative abundances. Significant differences in bacterial assemblages were assessed by permutational multivariate analysis of variance (PERMANOVA) with wUniFrac distances and the explanatory variables host species and depth (*i.e*., vegan::adonis) (Oksanen et al. 2017). Both overall (*i.e., O. annularis* and *O. franksi*) and species-specific models (*i.e.*, *O. annularis* or *O. franksi*) were tested. Heatmaps of OTU abundances were created using the phyloseq::plot_heatmap function (McMurdie and Holmes 2013). Within-category microbiome variability (*i.e.*, wUniFrac distance) was calculated in QIIME using the make_distance_boxplots function, which also assesses significant differences in microbiome variability among categories via pairwise, nonparametric t-tests (1000 Monte Carlo permutations) with Bonferroni correction. To test for significant differences in OTU abundances across host species and depths, we employed negative binomial modelling using DESeq2 (McMurdie and Holmes 2013; Love et al. 2014). Both the overall (*i.e.*, *O. annularis* and *O. franksi*) and species-specific models (*i.e.*, *O. annularis* or *O. franksi*) were tested. P-values for the significance of contrasts were generated based on Wald statistics, and false discovery rates were calculated using the Benjamini–Hochberg procedure.

### Microalgal communities

#### Internal Transcribed Spacer 2 rRNA (ITS2) amplicon library preparation, sequencing, and initial processing

To quantify differences in dinoflagellate communities across species and depths, we used a two-stage amplicon PCR on the same DNA that was extracted and used for the 16S amplification. We amplified the Internal Transcribed Spacer 2 (ITS2) rRNA marker gene commonly used for identification of Symbiodiniaceae (Hume et al. 2019). The primers used to quantify differences in the symbiotic algal communities were modified versions of the ITS-DINO forward (5’-ACA CTG ACG ACA TGG TTC TAC A<u>GT GAA TTG CAG AAC TCC GTG</u>-3’) and ITS2Rev2 (5’-TAC GGT AGC AGA GAC TTG GTC TCC T<u>CC GCT TAC TTA TAT GCT T-3</u>’) (Stat et al. 2009) that include the universal primer sequences required for Illumina MiSeq amplicon sequencing, namely common sequence 1 (CS1) and common sequence 2 (CS2). The PCR amplification was structured as follows: 2 min of denaturation at 94 °C; 35 cycles of 45 s at 94 °C, 60 s at 55 °C, and 90 s at 68 °C; then finally 7 min at 68 °C. Once the PCR reactions were finished, samples were held at 4 °C before sequencing. Samples were sequenced using the Illumina MiniSeq platform at the DNA Services Facility at the University of Chicago, Illinois. Sequences were submitted to SymPortal for processing and quality checks (Hume et al. 2019). Quality checking was performed using mothur (Schloss et al. 2009), followed by taxonomic identification using blastn. The SymPortal pipeline then subdivides sequences into genus groupings and identified type profiles, referred to as defining intragenomic sequence variants (DIVs). Type profiles were only identified if a variant contained more than 200 sequences, and the sequences were subsequently named based on whether they had been used in the definition of the DIVs. The resulting absolute and relative count tables were imported into R version 3.5.2 (R Core Team 2015) for downstream analyses and figure generation using the phyloseq (McMurdie and Holmes 2013), vegan (Oksanen et al. 2017), microbiome (Lahti and Shetty 2017), and ggplot2 (Wickham 2009) packages.

#### β-diversity group significance testing

To compare dinoflagellate communities across samples, we constructed Bray-Curtis and Jaccard dissimilarity matrices using absolute abundances. Significant differences in bacterial communities between sample types were assessed by PERMANOVA with Bray-Curtis and Jaccard distances and explanatory variables including host species, season, and depth using the adonis function from the vegan package (Oksanen et al. 2017). We tested overall models that encompassed both species as well as species-specific models.

## RESULTS

### Temperature and irradiance are higher and more variable in shallow environments

The *K*_d_ near the transplant sites was 0.40 m^-1^, indicating that corals from the shallow (3.5 m) and deep (9.5 m) sites receive respectively ~25% and ~2% sea surface irradiance. Across the vertical distribution range of each species (**Fig 1a**), it is estimated that the light intensity varies between 18% and 62% sea surface irradiance for *O. annularis* and between 5% and 33% for *O. franksi.* Relative light levels recorded by data loggers indicated that the light exposure was nearly 5 times more variable in shallow water than in deep water. Daily temperatures were significantly higher in the shallow site (28.85 ± 0.96 °C, mean ± s.d.) than in the deep site (28.46 ± 0.88 °C; *t*-value = 3.92, *p* < 0.001; **Fig. 1b**). However, based on the scaling quotient of temperature (Q_10_) of *Orbicella* spp. (Scheufen et al. 2017), it is estimated that the metabolic rate variation due to differences in temperature among sites is negligible (~5%).

**Figure 1.**
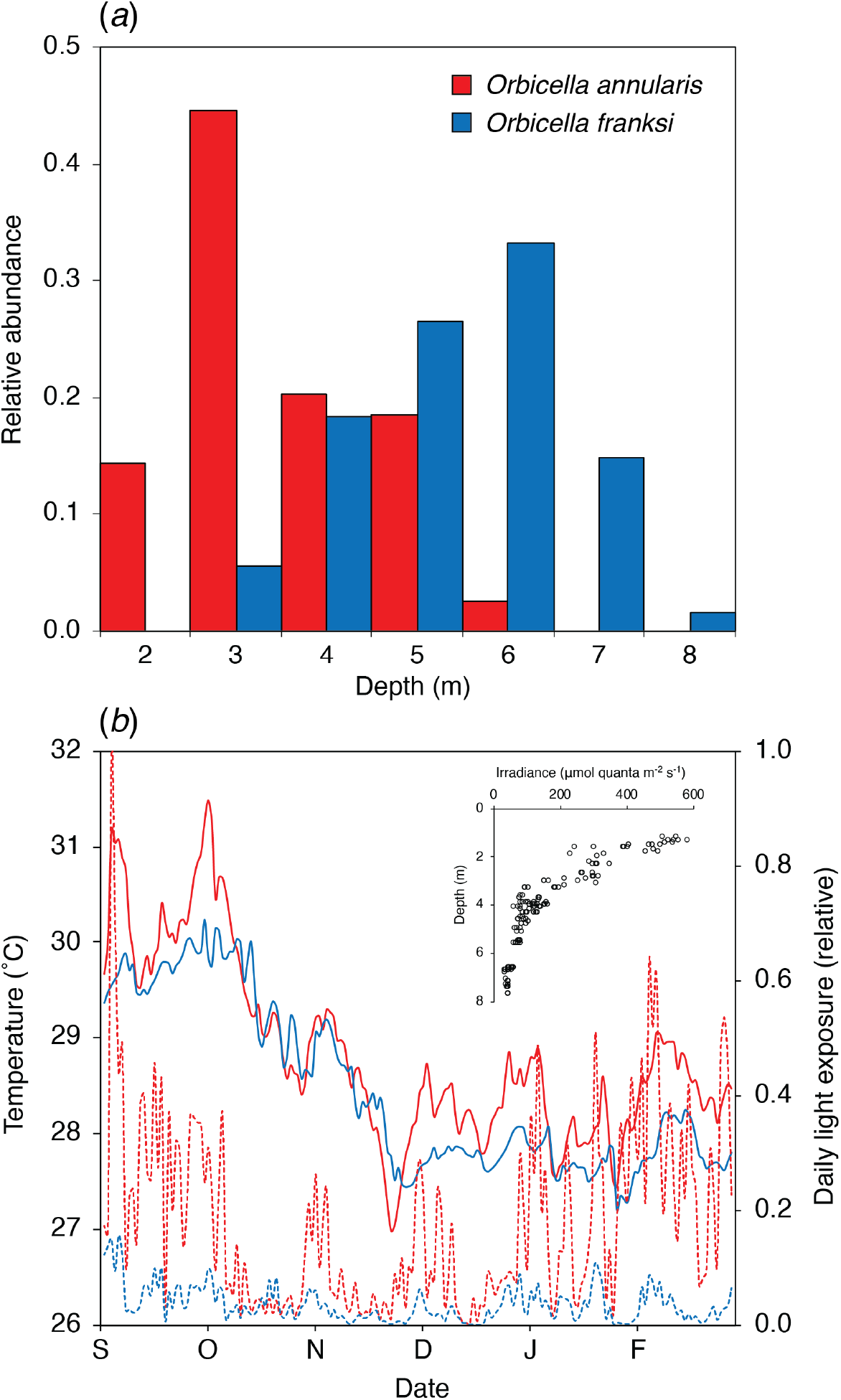
(*a*) Vertical distribution of *O. annularis* and *O. franksi* around the transplant sites in Bocas del Toro, Panamá, previously established as part of the long-term monitoring of coral spawning (Levitan et al. 2011). (*b*) Variation of the mean daily temperature (continuous lines) and relative light exposure (discontinuous lines) at the shallow (red) and deep (blue) transplant sites. The inset shows the light intensity variation across depths used to calculate the local *K*_d_.

### *O. annularis* experiences greater mortality in deep environments

Transplantation of *O. annularis* S-D (Δ_depth_ = 6 m) resulted in 26% mortality (Fisher exact test: *p* = 0.003) and was significantly higher than that of *O. franksi* colonies transplanted D-D (4% mortality, Fisher exact test: *p* = 0.04). *O. franksi* therefore has an advantage of 26% over *O. annularis* in deep habitats. In contrast, *O. franksi* when transplanted D-S did remarkably well with only 2% mortality (Fisher exact test: *p* = 0.63). Mortality of the two species was not significantly different (0% mortality, Fisher exact test: *p* = 0.60), suggesting that *O. franksi* in shallow areas has no perceivable short-term (< six months) disadvantage relative to *O. annularis* (Fisher exact test: *p* = 0.60).

### Photoacclimation of *O. annularis* is insufficient to compensate for reduced light

Symbionts of *O. annularis* exhibited a significant increase in *F*_v_/*F*_m_ when transplanted S-D (0.622 ± 0.034) relative to corals transplanted S-S (0.541 ± 0.007) (*t*-value = −6.25, *p* < 0.01). On the contrary, symbionts of *O. franksi* transplanted D-S experienced a reduction in *F*_v_/*F*_m_ (0.470 ± 0.052) relative to D-D transplants (0.630 ± 0.020; *t-*value = 0.55, *p* < 0.01). Transplantation of *O. annularis* S-D induced a significant reduction in *Q*_m_ (0.008 ± 0.076), relative to S-S transplantation (0.216 ± 0.163; *t*-value = 3.67, *p* < 0.01) (**Fig. *2a***); while *O. franksi* exhibited a significant increase in *Q*_m_ (0.226 ± 0.156) when transplanted D-S, relative to D-D transplants (0.073 ± 0.056; *t*-value = 3.26, *p* < 0.01) (**Fig. 2*b***).

**Figure 2.**
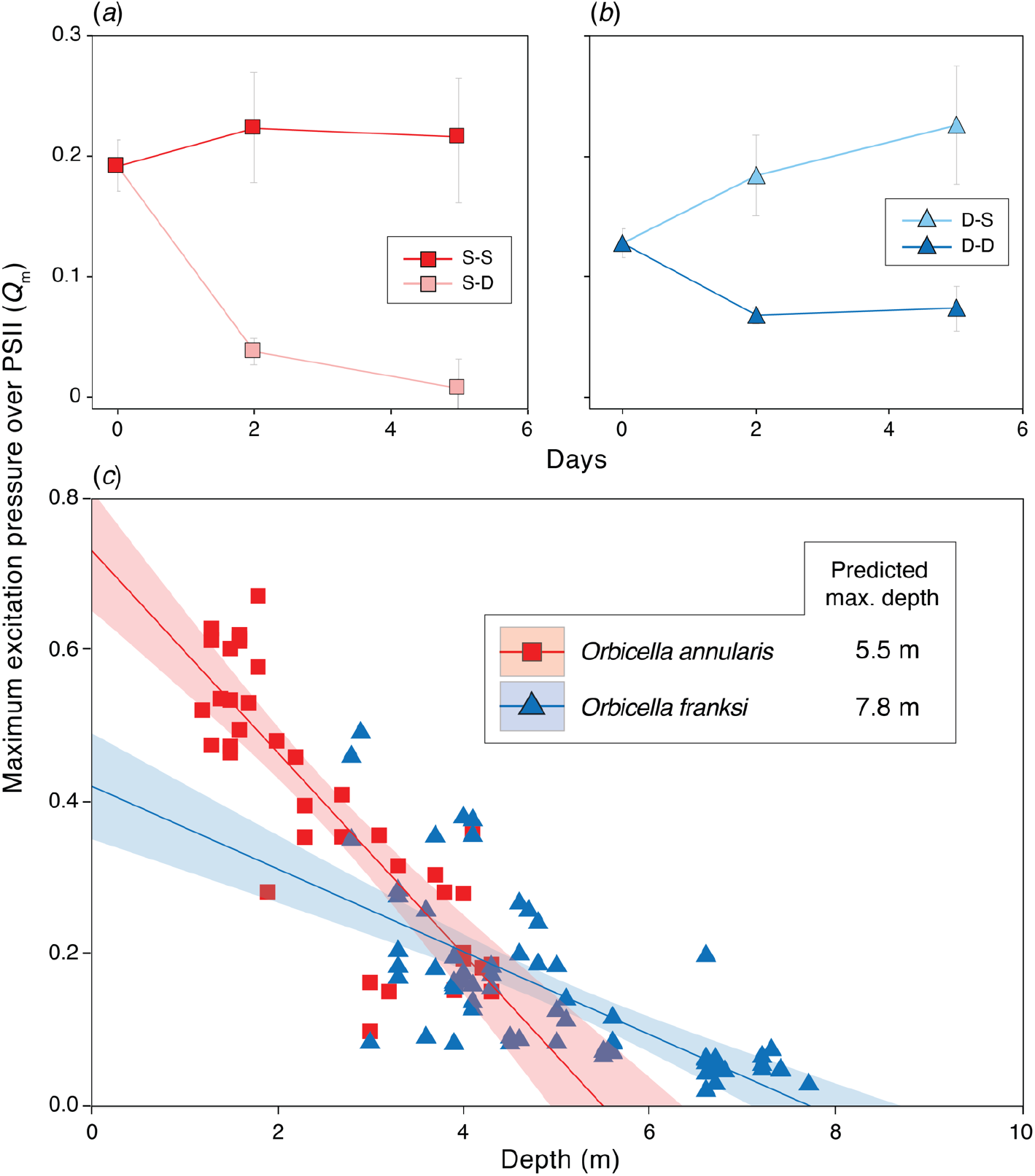
Photoacclimation responses of *Orbicella* spp. across depths. Maximum excitation pressure over PS II (*Q*_m_) is shown pre- and post-transplantation for *O. annularis* (*a*) and *O. franksi* (*b*). Values obtained in *O. annularis* transplanted S-S are shown in dark red while those transplanted S-D in pink. Values from *O. franksi* transplanted D-D are shown in dark blue while those transplanted D-S in light blue. (*c*) *Q*_m_ variation in *O. annularis* (red) and *O. franksi* (blue) along a depth gradient. A linear model was used to fit the data and predict the maximum potential depth limit described by *Q*_m_ for *O. annularis* [*Q*_m_ = 0.735 – 0.133*depth; *R*^2^=0.71, *p*<0.001] and *O. franksi* [*Q*_m_ = 0.422 - 0.054*depth; *R*^2^=0.50, *p*<0.001]. Clear lines represent 95% confidence intervals.

Estimations of *Q*_m_ on coral colonies along the vertical distribution of each species ranged from 0.099 to 0.673 in *O. annularis* and from 0.020 to 0.492 in *O. franksi* (**Fig. 2*c***). We found a significant species by depth interaction (F_(1,102)_=28.78, *p*<0.001), indicating that the slope of the regression model describing the relationship between *Q*_m_ and depth was significantly different between species, being more than twice as pronounced in *O. annularis* (m=-0.13; *R*^2^=0.71, *p*<0.001) than in *O. franksi* (m=-0.05; *R*^2^=0.50, *p*<0.001). The linear regression of *Q*_m_ with depth indicated that the potential depth limit described by the bioenergetics of the coral-algae symbiosis (*i.e.*, where *Q*_m_ reaches the minimum theoretical value of 0) is 5.5. m for *O. annularis* and 7.8 m for *O. franksi* (**Fig. 2c**), which nearly coincide with the observed lower limit of distribution of both species in the study area (**Fig. 1a**).

### Changes in depth produces a major shift in *O. annularis* microbiome

After quality control, sequencing resulted in a total of 577,930 microbial reads (per sample median: 5,758; per sample mean: 9,173) partitioned across 14,274 unique OTUs. Overall, coral-associated prokaryote communities were significantly structured according to depth (*p* = 0.001), but not host species (*p* = 0.12) or depth by species interaction (*p* = 0.86; PERMANOVA on weighted UniFrac; **Fig. 3**). The change across depths is mainly driven by *O. annularis* (*p* = 0.01, **Fig. 3**). The strong response of *O. annularis* microbiomes to changes in depth can be visualized in differential patterns of OTU abundance among depths (**Fig. *3a***).

**Figure 3.**
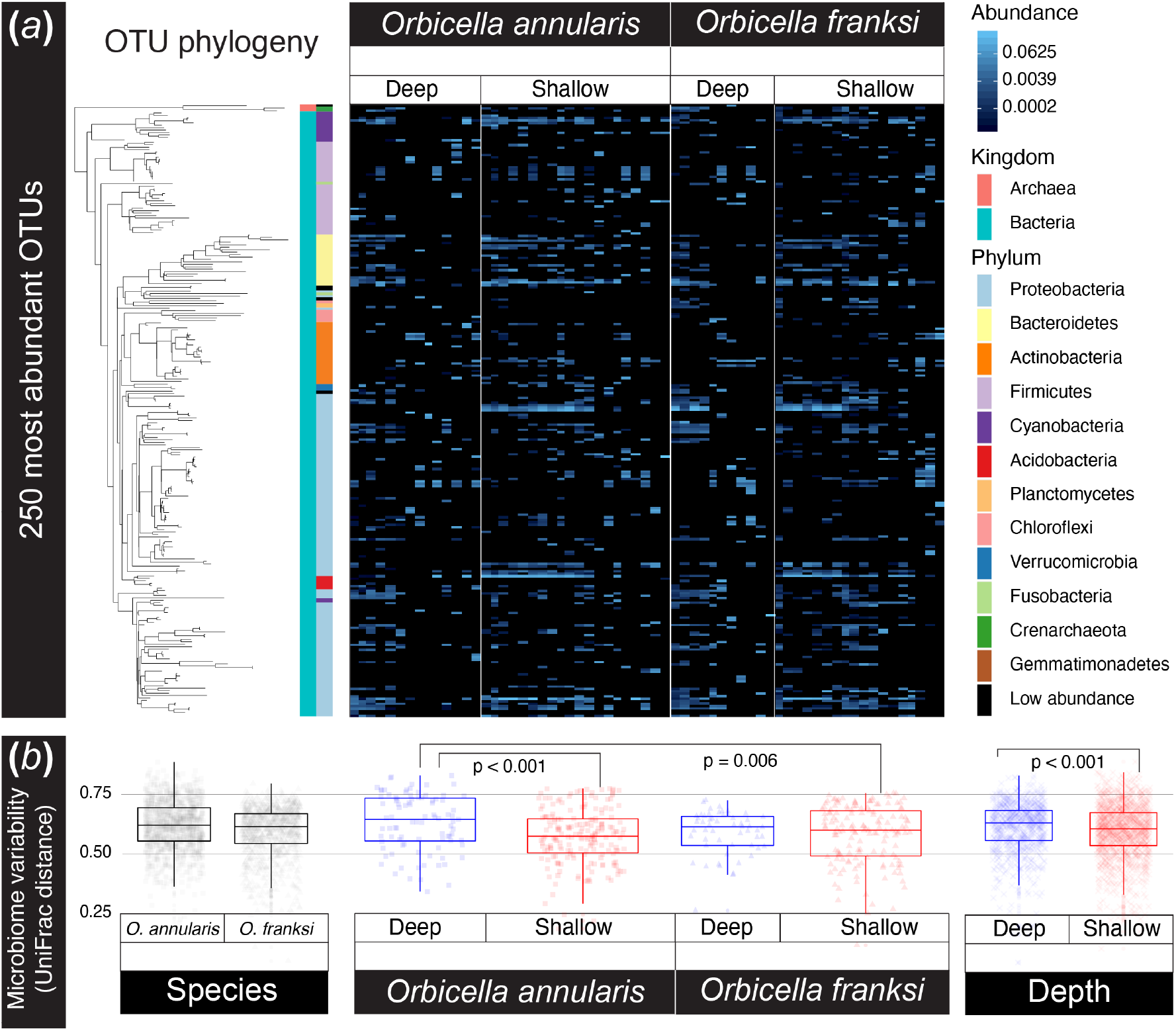
*O. annularis* microbiomes vary across timepoints and depths while *O. franksi* communities remain consistent. (*a*) Relative abundances of the 250 most common OTUs reveal distinct patterns among *O. annularis* microbiomes at the two transplant depths while *O. franksi* abundance patterns remain largely consistent across treatments. Each column in the heatmap represents an individual microbiome sample and phylogenetic relationships among OTUs are shown on the left (FastTree maximum-likelihood tree). (*b*) Microbiome variability (*i.e.,* weighted UniFrac distances) was greatest in *O. annularis* corals transplanted to deep waters. Microbiome variability was higher in corals in deep waters than in shallow.

Ten bacterial taxa were significantly enriched in shallow-water samples. OTUs enriched in shallow-water coral microbiomes are from the bacterial Orders Acidimicrobiales (1 OTU), Alteromonadales (1), Kiloniellales (2), Lactobacillales (1), Neisseriales (1), Oceanospirillales (3), and Synechococcales (1). The mean log2 fold change for enriched OTUs was 5.6.

Microbiome variability did not differ significantly between species with *O. annularis* (0.592 ± 0.008; mean UniFrac distance ± standard error) and *O. franksi* fragments (0.582 ± 0.008) (p_adj_ = 0.358). In contrast, microbiome variability differed significantly between depths, being greatest in *O. annularis* transplanted S-D (0.631 ± 0.013; mean UniFrac distance ± standard error) and significantly higher than *O. annularis* transplanted S-S (0.574 ± 0.008; p_adj_ < 0.001) or *O. franksi* transplanted D-S (0.580 ± 0.010; p_adj_ = 0.006) (**Fig. 3*b***). The larger microbiome variability in *O. annularis* transplanted deep is consistent with higher mortality and limited photoacclimation potential.

### Symbiodiniaceae communities vary across species

Algal communities of *O. annularis* were significantly different from those of *O. franksi* regardless of the depth to which they were transplanted to (*p* < 0.05; pairwise PERMANOVA on a Bray-Curtis matrix). Symbiodiniaceae genotypes belonging to the genus *Symbiodinium* (ITS2 type A3) and *Cladocopium* (C3an, C3an/C3, C7, and C7f) occurred in both coral species, although *Cladocopium* genotypes were more abundant in *O. franksi.* Genotypes from the genus *Breviolum* (B1 and B1/B1t) were detected in high abundances in *O. annularis,* and in many colonies from the shallow site (40% of them) were the only dominant symbiont. Only one *O. franksi* colony transplanted D-S hosted a *Breviolum* (B1) population. Genotypes belonging to the genus *Durusdinium* (D1, D1bl, D4, and D4c) were detected only in *O. franksi* transplanted D-D (**Fig. 4**). Neither *O. annularis* nor *O. franksi* Symbiodiniaceae communities were significantly different when transplanted to a different depth (*p* > 0.1; pairwise PERMANOVA on a Bray-Curtis matrix).

**Figure 4.**
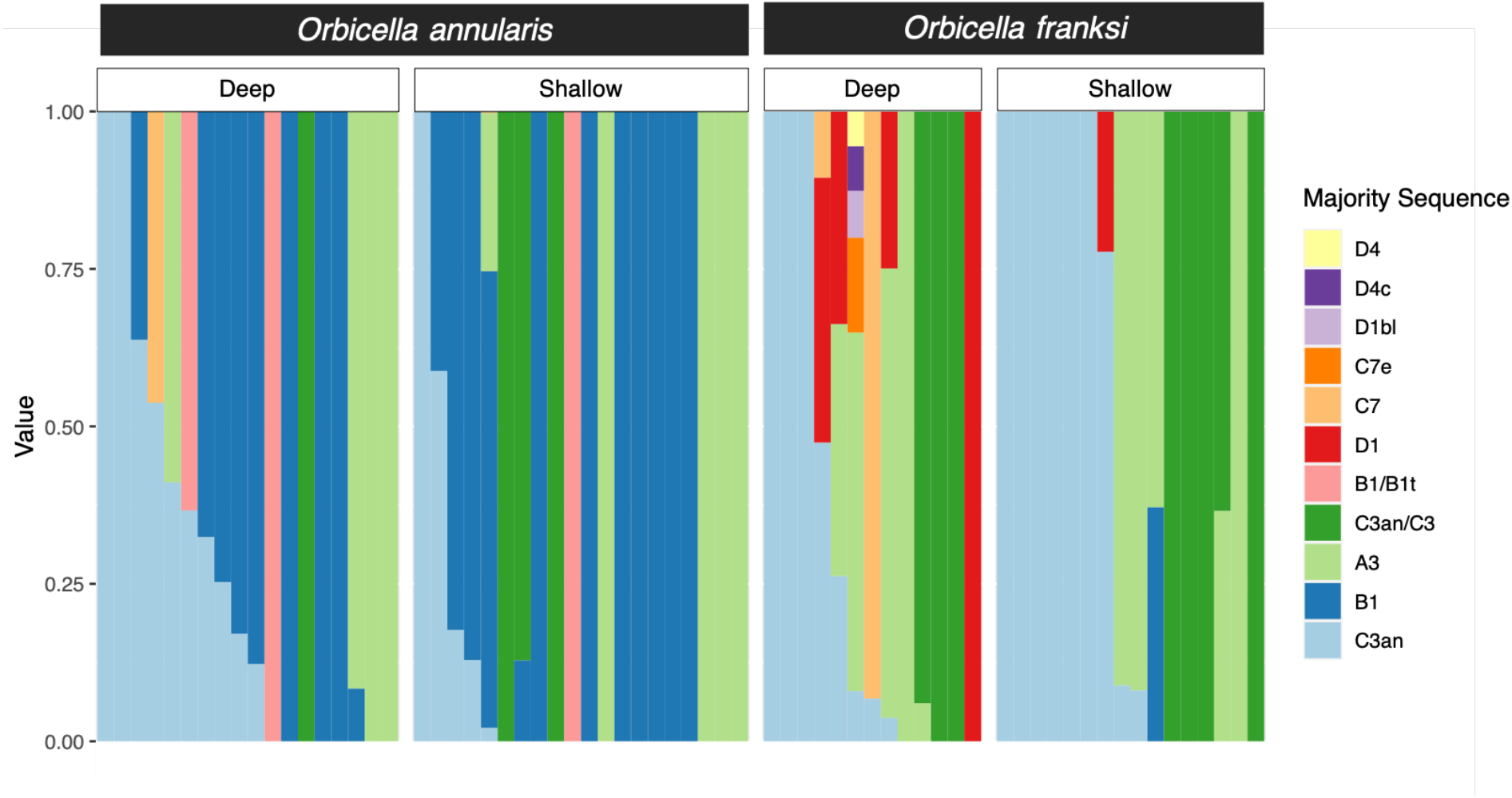
Relative abundance bar plot of Symbiodiniaceae ITS2 profiles identified in *Orbicella* spp. by Symportal (Hume et al. 2019). Variation in Symbiodiniaceae types is shown by species as well as by depth.

## DISCUSSION

Our study demonstrates that despite being genetically close (Levitan et al. 2011), *O. annularis* and *O. franksi* have diverged physiologically and occupy distinct light environments in part due to the variation in their associated microbiotas (Symbiodiniaceae and bacterial communities). Following transplantation to deep habitats, *O. annularis* experiences a limited photoacclimation potential and disruption of the photosynthetic performance of its algal symbionts, consistent with increased mortality and significant microbiome community shifts with increased variability. By contrast, *O. franksi* maintained a robust physiological performance, a resilient microbiome composition with no significant community shifts or increased variability, and low mortality at both depths. Our study suggests that *O. annularis* is adapted to shallow environments characterized by a higher and more variable temperature and light regimes, while *O. franksi* is physiologically able to live in both shallow and deep habitats. The niches of these sibling species have diverged, and a large component of the niche separation seems to be related to variations in the photoacclimation capabilities and the microbial community of each species. The absence of *O. franksi* in shallow areas may be related to other ecological aspects such as slow growth in an area of intense space competition, a restricted morphological plasticity for regulating the light capture (see below) and/or non-random settlement of recruits across depths (*i.e.,* larval habitat choice).

### The vertical distribution couples with the photoacclimation capabilities of each species

The vertical distribution of *O. annularis* and *O. franksi* is compressed toward shallower depths in Bocas del Toro compared to other clear-water sites in the Caribbean (*e.g*., Curaçao (Van Veghel 1994) and Belize (Pandolfi and Budd 2008)). The vertical habitat compression in both species is consistent with the *K*_d_ measured in Bocas del Toro (0.40 m^-1^), which is notably higher than in clear-water sites (0.06 m^-1^ in Curaçao and 0.08 m^-1^ in Belize (Banaszak et al. 1998; Vermeij and Bak 2002)) and reflects the effect of the heavy rainfall patterns and runoff in the region on the optical properties of the water column (Kaufmann and Thompson 2005). This vertical habitat compression is consistent with other coral reefs exposed to water turbidity (Morgan et al. 2020; López-Londoño et al. 2021) and suggest that the light penetration into the water column associated with the local *K*_d_ is a determinant factor for the vertical zonation of *Orbicella* spp. Despite local differences in the vertical distribution ranges, *O. annularis* consistently occupies well-lit shallow areas of reefs where the potential for increased photosynthesis and calcification rates drives a steep competition for space with other corals. In contrast, *O. franksi* consistently dominates deeper reef areas characterized by low-light conditions and reduced coral-growth rates (Cohen and Dubinsky 2015).

Our findings indicate that *O. annularis* experiences an almost complete loss of photosynthetic activity when transplanted deep. *O. annularis* fragments photoacclimate to low-light conditions by increasing the light energy conversion efficiency (*i.e*, increase in *F*_v_/*F*_m_) (Hoegh-Guldberg and Jones 1999; Gorbunov et al. 2001). However, the extremely low values of *Q*_m_ reflect a trivial photosynthetic contribution of *O. annularis* symbionts to the host metabolism due to light-limited photosynthesis (Iglesias-Prieto et al. 2004), suggesting that the photoacclimation potential is insufficient to compensate for the low-light conditions of deep environments. Photoacclimation of *O. franksi* fragments transplanted to the shallow environment resulted in an increased fraction of photo-inactivated PSII reaction centers and capacity for thermal dissipation of excessive light energy absorbed (Hoegh-Guldberg and Jones 1999; Gorbunov et al. 2001). But in contrast to *O. annularis,* the estimated *Q*_m_ in *O. franksi* do not indicate the occurrence of chronic photoinhibition in the shallow environment nor light-limitation in the deep environment, suggesting that *O. franksi* can maintain a more robust physiological performance across depths. The photoacclimation responses of both species in the transplant experiment were consistent with the rates of change in *Q*_m_ across their vertical distribution range, which collectively suggest that the symbiotic algae of *O. annularis* are more sensitive to changes in light intensity with depth than symbionts of *O. franksi*.

Colony morphology can help modulate the light capture and photosynthetic energy acquisition along the vertical distribution range of corals (Hoogenboom et al. 2008; Kaniewska et al. 2011). The dominance of *O. annularis* in shallow habitats correlates with its faster vertical growth among *Orbicella* species (Weil and Knowlton 1994). Its morphology (typically columnar) helps regulate the distribution of light energy for symbiotic algae across the colony surface, representing an advantageous strategy in high-light environments because it reduces the coral tissue area subjected to excessive irradiance (Kaniewska et al. 2011). When transplanted deep, this morphology may lead to acute light energy limitation which, in combination with the insufficient acclimation potential to compensate for low-light, can lead to negative energetic balances for the whole colony and eventual death. *O. franksi,* on the other hand, produce plate-like colonies to maximize light capture in deep environments. When transplanted to shallow well-lit environments, despite a potential for successful photoacclimation as indicated by our results, the plate-like morphology limits the capacity to regulate the internal light climate and allows very slow vertical growth. This slow growth makes *O. franksi* a poor competitor, likely explaining why this species is rare in shallow areas. Alternatively, and not mutually exclusive, their larvae may preferentially settle in low light environments. In fact, adaptation and strong selection across depths, may promote the evolution of habitat choice.

### Host species drive symbiont communities

Species-specific associations with algal symbionts with contrasting photoacclimation capabilities may be a key axis of differentiation between *O. annularis* and *O. franksi.* Despite the higher and more variable temperature and light intensity in shallow areas, which are known conditions that promote the association with *Durusdinium trenchii* in other corals (LaJeunesse et al. 2009), this dinoflagellate was not detected in *O. annularis* colonies. Surprisingly, this thermotolerant symbiont (ITS type D1/D1bl) was found in nearly 20% of *O. franksi* colonies from the deep environment. The increased abundance of *D. trenchii* in *O. franksi* may be related with the runoff impacts in the water column (*e.g.,* sedimentation and nutrient enrichment), a reduction in light penetration, and the mechanisms by which the coral-algae symbiosis interact with these environmental conditions (Garren et al. 2006). The prevalence of *Breviolum* genotypes in *O. annularis* and *Cladocopium* genotypes in *O. franksi,* both in the shallow and deep transplant sites, is consistent with previous reports (LaJeunesse 2002; Garren et al. 2006) and may indicate the formation of stable associations explained by the photoacclimative capabilities of dinoflagellates and the variability of physical factors within the vertical distribution range of each coral species (LaJeunesse 2002; Iglesias-Prieto et al. 2004). The ITS2 analysis has a low resolution to differentiate linages within the same genus in symbiotic algal communities (LaJeunesse and Thornhill 2011; Stat et al. 2011). It is possible that complementary analysis with other molecular markers improves the phylogenetic resolution of Symbiodiniaceae (*i.e.,* species or population level), detecting differences in cryptic species/populations of *Cladocopium* spp. or *Breviolum* spp. uniquely associated with each *Orbicella* species like in other depth-segregated anthozoans (Prada et al. 2014; Pochon et al. 2015).

### Microbiome communities vary across depths and are enriched in shallow habitats

Several Endozoicomonas OTUs were significantly enriched in shallow habitats. Endozoicimonaceae are diverse gammaproteobacterial symbionts of numerous marine hosts at varying depths and with a wide global distribution (Neave et al. 2016). Members of this group are found in abundance in the tissues of coral species and are considered to be true symbionts of corals which may provide a beneficial function (Bayer et al. 2013; Pantos et al. 2015). Although their function within the coral host is not entirely clear; proposed benefits include nutrient acquisition, microbiome structuring and roles in coral health.

Members of the family Alteromonadaceae and the order Acidimicrobiales were also enriched in shallow areas. Alteromonadaceae belong to a diverse group of heterotrophic gammaproteobacteria known to associate with marine hosts and nutrient rich environments. Members of this group tolerate relatively high temperatures and have been used in coral probiotic studies as coral-associated bacteria capable of scavenging free radicals (Dungan et al. 2020), and therefore could provide similar benefits in shallow, high-light environments. Similarly, Acidimicrobiales are known to be planktonic free-living photo-heterotrophs found in both temporal and tropical photic zones (Angly et al. 2016) and are associated with DOM in marine environments (Osterholz et al. 2018).

Finally, corals in shallow areas were also enriched for *Alloiococcus* and *Synechococcus. Alloiococcus* belongs to the group of gram-positive lactic acid bacteria, which are recognized for producing bacterial growth inhibitors that function to deter invading bacteria in their hosts (Ringø et al. 2018). *Synechococcus* is a photoautotrophic cyanobacterium found in surface waters harbouring abundant light. Both corals and their symbiotic algae are known to actively feed on *Synechococcus* (Jeong et al. 2012; McNally et al. 2017) which is often found as a member of the coral surface mucus microbiome (Marchioro et al. 2020). As a food for corals, it has been suggested that nitrogen-rich *Synechococcus* cells may increase bleaching recovery and coral health (Meunier et al. 2019).

There is a continuing debate as to the relative role of coral host vs. environment in shaping coral microbiomes. This study demonstrates that the responsiveness of coral microbiomes to environmental conditions differs significantly even among very closely related coral species. These differences in microbiome shifts may be related to the resilience of the coral host and its associated algal community to a particular habitat. Pantos et al. (Pantos et al. 2015) found that environment is the major driver of microbiome structure in *Seriatopora hysterix,* not host genotype or Symbiodiniaceae strain. Our results do not contradict this finding but suggest that responsiveness to environmental conditions can differ significantly even among very closely related coral taxa.

### Implications for coral reef conservation

A key finding in our study with implications for coral restoration is the increased mortality of *O. annularis* when transplanted to low-light environments. We suggest that to enhance survivorship during restoration, the particular light environment of source populations should be similar to the transplant sites. In this study, due to the high vertical attenuation of light (*K*_d_ = 0.40 m^-1^), a 6 m increase in depth resulted in an order of magnitude reduction in irradiance and increased mortality of *O. annularis* by 26%. In a clear-water site (*e.g*., *K*_d_ = 0.06 m^-1^), this response would be expected to occur with an increase in depth of ~40 m. Giving the expensive nature of coral restoration, equating the light environment of donor and transplant sites will likely increase yield and decrease costs. Minimally, our approach can be used to estimate the maximum theoretical depth for each species in a given location with certain water optical quality, thereby providing guidance when choosing the location and depth for coral transplantation.

The second aspect of our findings is related to microbiome composition in different habitats across reefs. Shallow water reefs are areas of high stress with strong variations in light, temperature and salinity, strong changes in water motion and sediment transport, and more ecological variability. Our study suggests that the microbiome of shallow water specialists like *O. annularis* is fine-tuned to this environment and a reduction in the light field can cascade into drastic changes to host-associated microbial community composition. Increases in temperature as a result of climate change has affected primarily shallow water corals (Bridge et al. 2013; Hughes et al. 2018), suggesting that instability in the coral microbiomes of shallow-water corals will increase, likely accelerating coral decline of these reef areas.

Lastly, subtle differences in the water optical conditions can result in changes in the underwater light environment and the vertical distribution of coral species. Most coral reefs around the globe are currently threatened by the direct effects of sediments, pollutants and nutrients associated with coastal development and terrestrial runoff (Carlson et al. 2019). These conditions affect the water optical quality and, as a consequence, the light climate of corals and the survivorship of species at different depths. Although previous studies have suggested that deep-water species are more sensitive to changes in water optical conditions (Vermeij and Bak 2002), our results suggest that at least some shallow-water specialists, like *O. annularis,* can be extremely vulnerable to these changes as their physiology/morphology is specialized for high light habitats. As the degradation of water optical properties in coral reefs continue, shallow-water specialists, which are typically major reef-building species, will likely become rare, shifting the structural and functional integrity of reefs.

## CONCLUSION

Our study suggests that the sibling coral species, *O. annularis* and *O. franksi,* are adapted to distinctive light environments along depth gradients. The limited photoacclimation potential and less robust microbiome community restricts *O. annularis* to shallow, high-light environments. *O. franksi* is more versatile, but other ecological aspects such as slow growth in areas of intense space competition restricts the species to deep environments. These contrasting responses associated with the microbial communities highlight the importance of niche specialization in symbiotic corals for the maintenance of species diversity. Our study has implications on coral reef restoration efforts, providing guidance when choosing the location, depth and light environment for coral transplantation.

## ACKNOWLEDGMENTS

Arcadio Castillo Díaz, Gabriel Jácome and Plinio Góndola from the Smithsonian Tropical Research Institute assisted with field operations at the Bocas del Toro field station. Gaby Swain helped on sample processing. Dr. Benjamin Hume provided assistance on Symbiodiniaceae analysis through the SymPortal framework.

## FUNDING STATEMENT

This work was supported by NSF grants OCE 1442206 and OCE 1642311; Pennsylvania State University startup funds to MM and RI-P; and NOAA grant NA19NOS4820132. CP was funded by grants from NSF (OIA) 2032919 and USDA National Institute of Food and Agriculture (Hatch) 1017848.

## COMPETING INTERESTS

The authors declare that they have no competing interests.

## Notes

### Competing Interest Statement

The authors have declared no competing interest.

